# Cell freezing protocol optimized for ATAC-Seq on motor neurons derived from human induced pluripotent stem cells

**DOI:** 10.1101/036798

**Authors:** Pamela Milani, Renan Escalante-Chong, Brandon C. Shelley, Natasha L. Patel-Murray, Xiaofeng Xin, Miriam Adam, Berhan Mandefro, Dhruv Sareen, Clive N. Svendsen, Ernest Fraenkel

## Abstract

In recent years, the assay for transposase-accessible chromatin using sequencing (ATAC-Seq) has become a fundamental tool of epigenomic research. However, it has proven difficult to perform this technique on frozen samples because freezing cells before extracting nuclei impairs nuclear integrity and alters chromatin structure. We describe a protocol for freezing cells that is compatible with ATAC-Seq, producing results that compare well with those generated from fresh cells. We found that while flash-frozen samples are not suitable for ATAC-Seq, the assay is successful with slow-cooled cryopreserved samples. Using this method, we were able to isolate high quality, intact nuclei, and we verified that epigenetic results from fresh and cryopreserved samples agree quantitatively. We developed our protocol on a disease-relevant cell type, namely motor neurons differentiated from induced pluripotent stem cells from a patient affected by spinal muscular atrophy.

## Introduction

Since its establishment, the assay for transposase-accessible chromatin using sequencing (ATAC-Seq) has revolutionized the study of epigenetic^1,2^. This technique detects open-chromatin regions and maps transcription factor binding events genome-wide by means of direct *in vitro* transposition of native chromatin. Specifically, hyperactive Tn5 transposase is used to interrogate chromatin accessibility by inserting high-throughput DNA sequencing adapters into open genomic regions, which allows for the preferential amplification of DNA fragments located at sites of active chromatin. Because the DNA sites directly bound by DNA-binding proteins are protected from transposition, this approach enables the inference of transcription factor occupancy at the level of individual functional regulatory regions. Furthermore, ATAC-Seq can be utilized to decode nucleosome occupancy and positioning, by exploiting the fact that the Tn5 transposase cuts DNA with a periodicity of about 150-200bp, corresponding to the length of the DNA fragments wrapped around histones^3^. This periodicity is maintained up to six nucleosomes and provides information about the spatial organization of nucleosomes within accessible chromatin. ATAC-Seq signals thus allow for the delineation of finescale architectures of the regulatory framework by correlating occupancy patterns with other features, such as chromatin remodeling and global gene induction programs. Compared to other epigenetic methodologies, such as FAIRE-Seq and DNase-Seq, ATAC-Seq requires relatively few cells and is less time-consuming and labor-intensive. Therefore, it is suitable for work on precious samples, including differentiated cells derived from induced pluripotent stem cells (iPSCs), primary cell culture, and limited clinical specimens.

It is essential to preserve the native chromatin architecture and the original nucleosome distribution patterns for ATAC-Seq. Freezing samples prior to the purification of nuclei can be detrimental to nuclear integrity and chromatin structure, thus restricting the application of ATAC-Seq to freshly-isolated nuclei. This limits the use of ATAC-Seq on clinical samples, which are typically stored frozen, and represents a major logistical hurdle for long-distance collaborative projects, for which sample freezing is often inevitable.

In an attempt to overcome this drawback, we developed an optimized freezing protocol suitable for native chromatin-based assays. We tested two different freezing methods: flash-freezing and slow-cooling cryopreservation. Flash-freezing is a procedure in which the temperature of the sample is rapidly lowered using liquid nitrogen, dry ice or dry ice/ethanol slurry, in order to limit the formation of damaging ice crystals. Conversely, slow-cooling cryopreservation lowers the temperature of the sample gradually and makes use of cryoprotectants, such as dimethyl sulfoxide (DMSO), to prevent ice crystal nucleation and limit cell dehydration during freezing. Cryopreservation techniques are widely employed for cell banking purposes and are routinely used in assisted reproduction technologies^4,5^.

We tested the freezing techniques using disease-relevant cell types, namely motor neurons (iMNs) differentiated from human iPSCs, which were derived from the fibroblasts of a patient affected by spinal muscular atrophy (SMA). This disease is caused by homozygous loss of the *SMN1* gene and is characterized by the degeneration of lower motor neurons^6^.

We introduced a number of experimental quality control (QC) checkpoints and steps for data analysis to monitor the efficacy of the procedures and quantify potential alterations induced by cell freezing. The method we describe should be applicable in a wide variety of settings and expand the number and types of samples that can be studied using ATAC-Seq.

## Results and Discussion

### Description of experimental design and overview of the protocol

We generated ATAC-Seq data on fresh (F), flash-frozen (FF), and cryopreserved (C) iMNs by following the procedure outlined in Figure 1. Fresh and frozen neurons were derived from the same pool of cells and processed in parallel in order to estimate the effects of freezing on ATAC-Seq outcomes without any batch effect bias.

**Figure 1.**
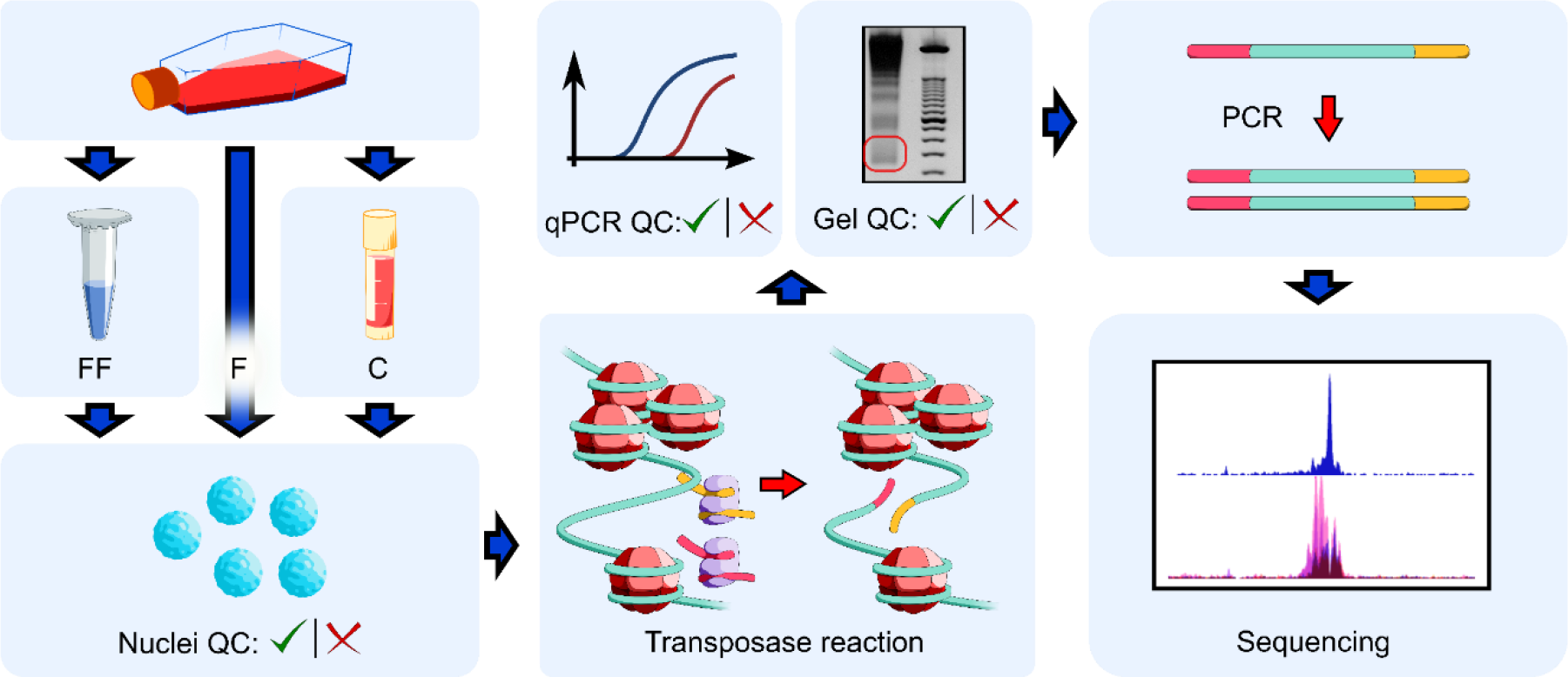
Outline of ATAC-Seq procedure using fresh, flash-frozen, and cryopreserved iPSC-derived motor neurons. The key experimental steps and quality control (QC) checkpoints are indicated, consisting of morphological evaluation of nuclei, agarose gel electrophoresis of libraries, and real-time qPCR to assess the enrichment of open-chromatin sites. (F = fresh, FF = flash-frozen, C = cryopreserved).

The ATAC-Seq protocol was adapted from Buenrostro et al^1,7^, with some modifications. Given that a successful ATAC-Seq experiment begins with the isolation of high-quality intact nuclei, we first introduced a quality control checkpoint consisting of the morphological evaluation of nuclei with either Trypan Blue or DAPI staining, followed by the accurate quantification of those nuclei using an automated cell counter. Precise counting of nuclei is important to ensure optimal tagmentation (the simultaneous fragmenting of the DNA and insertion of adapter sequences) and to limit the technical variability across samples. From a qualitative perspective, individual intact nuclei with a round or oval shape should be observed with no visible clumping. To exclude samples with severe degradation or over-tagmentation, we assessed the quality of the treated chromatin samples by gel electrophoresis, as described in Buenrostro et al.^1^; if the chromatin was intact and the transposase reaction was optimal, a DNA laddering pattern with a periodicity of about 200bp should be observed, corresponding to fragments of DNA that were originally protected by an integer number of nucleosomes (nucleosome phasing). Furthermore, we measured the enrichment of DNA accessible regions by performing realtime qPCR analysis using a known open-chromatin site as a positive control and a Tn5-insensitive site as a negative control. When assayed by real-time qPCR, high-quality ATAC-Seq samples should show at least a 10-fold enrichment of positive control sites compared to Tn5-insensitive sites. Finally, as we were principally interested in open-chromatin profiling and not in nucleosome positioning, we introduced a size-selection step to enrich for nucleosome-free fragments. This step increases the signal-to-noise ratio and improves the sensitivity of the methodology. After size-selection, libraries were PCR-amplified and submitted for single-end sequencing.

### ATAC-Seq on iPSC-derived motor neurons (iMNs): flash-frozen cells

We first performed ATAC-Seq on fresh and flash-frozen iMNs. Differentiated neuronal cells were generated as described in Methods. We performed immunocytochemistry experiments using antibodies against markers of mature motor neurons to test the efficiency of the differentiation protocol; we showed that patient-derived iPSCs were successfully differentiated into ISL1-and SMI32-positive motor neurons (Figure 2). Figure 3 shows ATAC-Seq outcomes from two representative samples. Nuclei from fresh cells passed quality control, while nuclei from flash-frozen neurons exhibited excessive clumping, likely caused by disruption of the nuclear envelope and consequent leakage of DNA (Figure 3A). After the transposase reaction, we assessed the quality of the resulting libraries by qualitative evaluation of agarose gel electrophoresis. The library from freshly-isolated nuclei displayed clear nucleosome phasing, while the library from flash-frozen neurons showed DNA smearing on the gel (Figure 3B). This result strongly indicates that loss of chromatin integrity occurred during flash-freezing. We proceeded with next-generation sequencing for one fresh and one flash-frozen sample and uploaded the data on the UCSC Genome Browser for manual inspection of tracks and local visualization of peaks (Figure 3C). As a negative control, we treated human naked DNA with the hyperactive Tn5 enzyme and sequenced this library alongside the ATAC-Seq samples. ATAC-Seq peaks from fresh neurons were sharp and overlap with H3K4me3 signals from ENCODE ChIP-Seq datasets. Using a MACS2 q-value threshold of 0.05, we obtained more than seventy thousand significant peaks using fresh cells. In contrast, the reads from flash-frozen cells were distributed evenly across the entire genome, similar to the results obtained with the negative control, and less than five hundred significant peaks were detected. These findings indicate that flash-freezing of iMNs is not suitable for ATAC-Seq.

**Figure 2.**
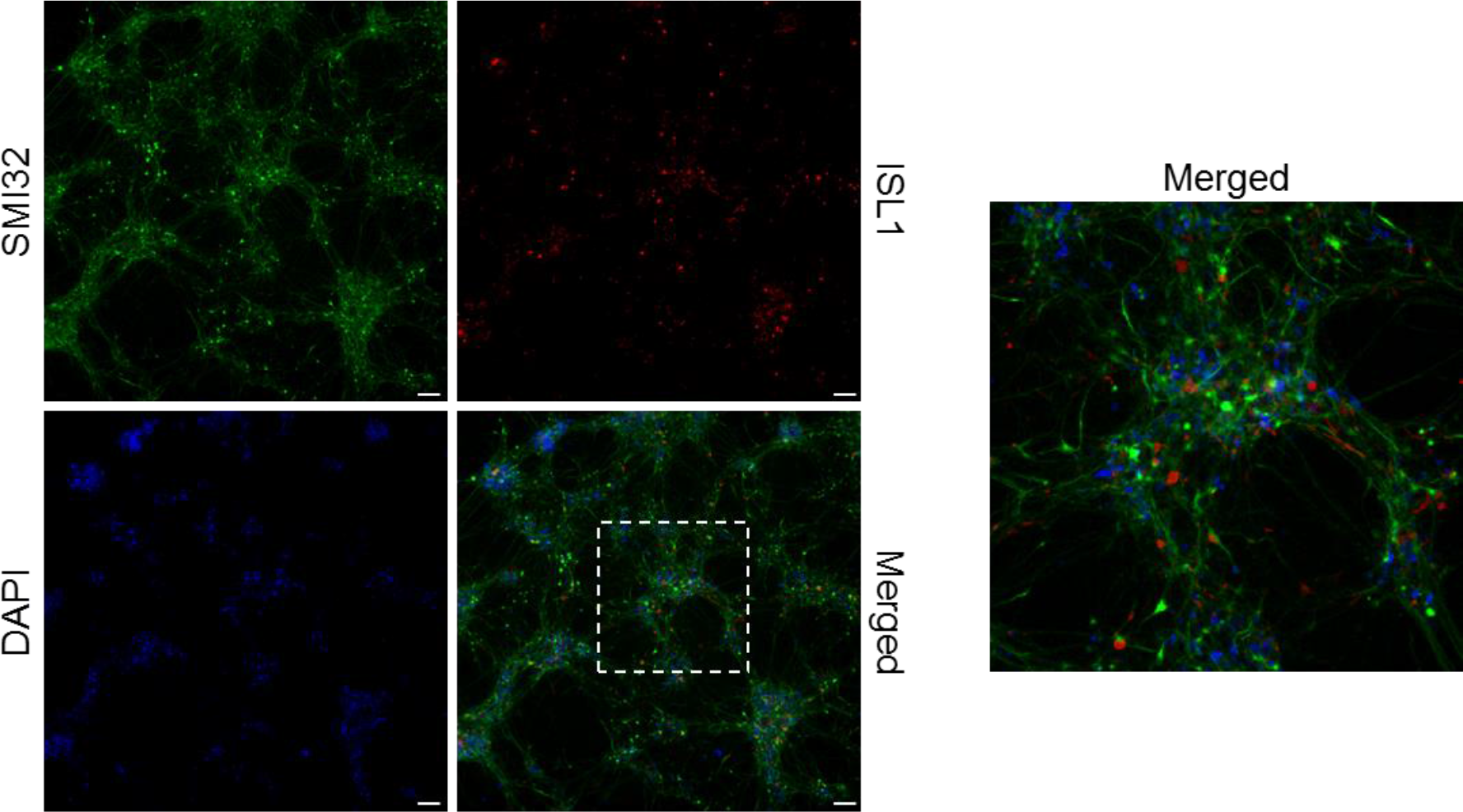
Fibroblast-derived iPSCs differentiate into SMI32-and ISL1-positive motor neurons. Differentiated cells were labeled to evaluate the immunoreactivity of SMI32 (green) and ISL1 (red) proteins, two markers of mature motor neurons. Nuclei were stained with DAPI. Motor neurons were imaged with 10x magnification. The image on the right represents a higher magnification of selected neurons. Scale bar = 75 μm.

**Figure 3.**
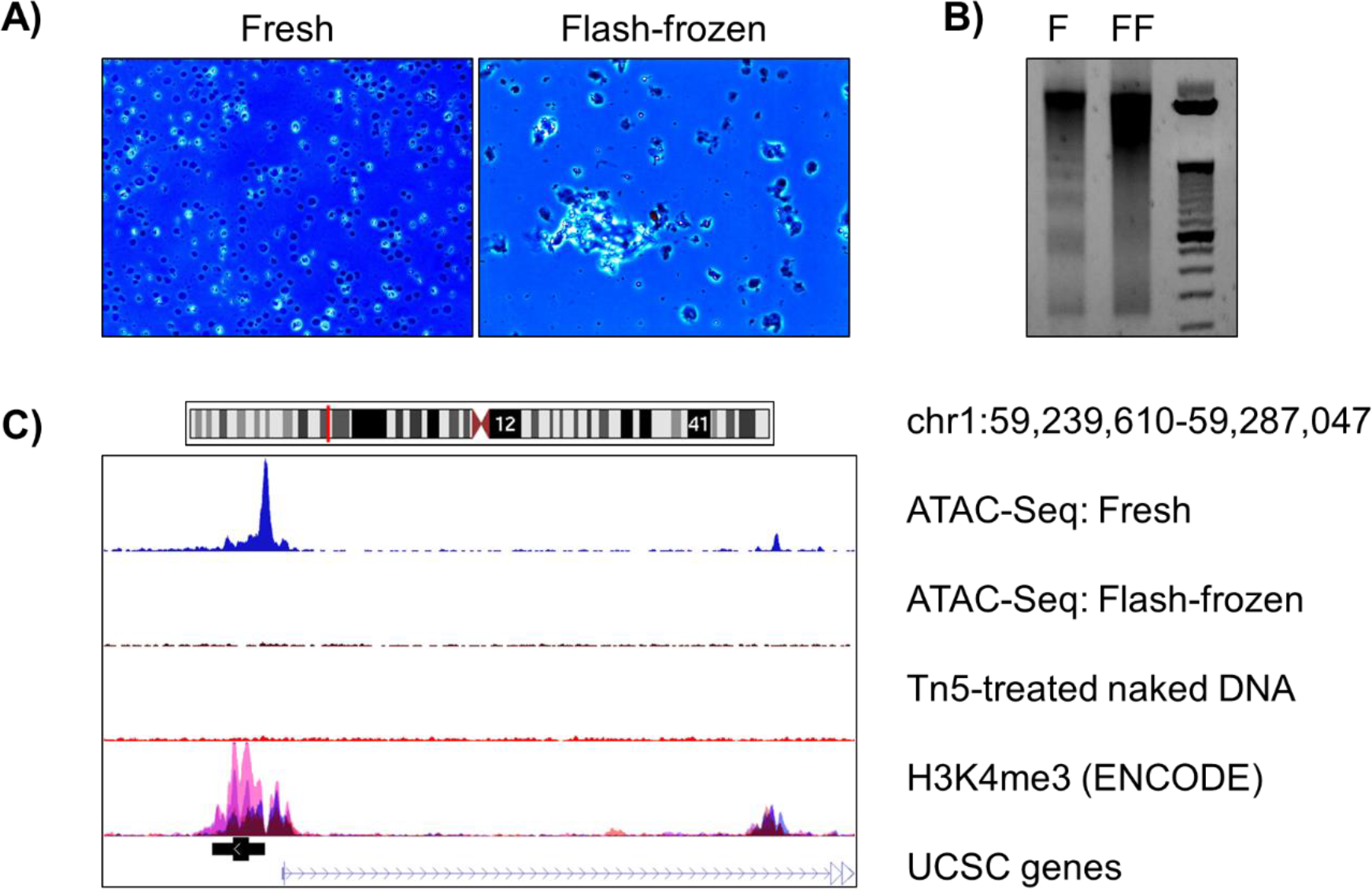
Representative results for ATAC-Seq carried out on fresh and flash-frozen cells. (A) Nuclear morphological evaluation: nuclei from fresh cells were of high quality, while excessive clumping was observed for nuclei from flash-frozen neurons. (B) Agarose gel electrophoresis of libraries: the nucleosome phasing pattern on the gel was not detected in flash-frozen samples, as opposed to fresh cells. (C) ATAC-Seq tracks uploaded on UCSC Genome Browser: while we detected sharp peaks for fresh samples, the reads from flash-frozen neurons were distributed noisily across the genome. (F = fresh, FF = flash-frozen).

### ATAC-Seq on iPSC-derived motor neurons (iMNs): cryopreserved cells

Next, we compared ATAC-Seq results from fresh and cryopreserved cells. Fresh iMNs were transferred to Cryostor media and slowly frozen, stored, and then thawed for processing (see Methods). As shown in Figure 4A, nuclei from the cryopreserved cells were of high quality and the nucleosome laddering was detected by gel electrophoresis (Figure 4B). Sequencing data from both fresh and cryopreserved samples showed sharp peaks and low background signal (Figure 4C). Furthermore, the qPCR enrichment of the positive control site *(GAPDH* gene promoter, Figure 5A top panel) over the Tn5-insensitive site (gene desert region, Figure 5A bottom panel) was high and comparable to that of fresh cells, as opposed to qPCR results from flash-frozen neurons, for which less than 10-fold enrichment was observed (Figure 5B). As in the case of fresh cells, we obtained more than seventy thousand significant peaks using cryopreserved samples (MACS2 q-value threshold = 0.05) (Table 1). These results reveal that slow-cooling cryopreservation is compatible with native chromatin-based epigenetic assays.

**Table 1.** Information about sequencing data. The numbers of total and aligned reads are indicated. Although the percentage of mitochondrial DNA (mtDNA) contamination is higher in cryopreserved samples when compared to fresh samples, the number of significant peaks mapping to nuclear DNA is similar across all samples. (F = fresh, C = cryopreserved).

| Sample | # of total reads | # of aligned reads | mtDNA (%) | # of significant peaks |
| --- | --- | --- | --- | --- |
| **F1** | 26,092,754 | 22,059,551 | 32.14 | 71,050 |
| **F2** | 30,730,456 | 25,950,925 | 27.97 | 70,073 |
| **F3** | 31,364,716 | 26,333,862 | 31.44 | 73,305 |
| **C1** | 28,201,642 | 24,577,487 | 45.33 | 73,973 |
| **C2** | 29,964,823 | 25,900,608 | 49.61 | 72,512 |
| **C3** | 28,018,248 | 23,660,777 | 47.69 | 70,547 |

**Figure 4.**
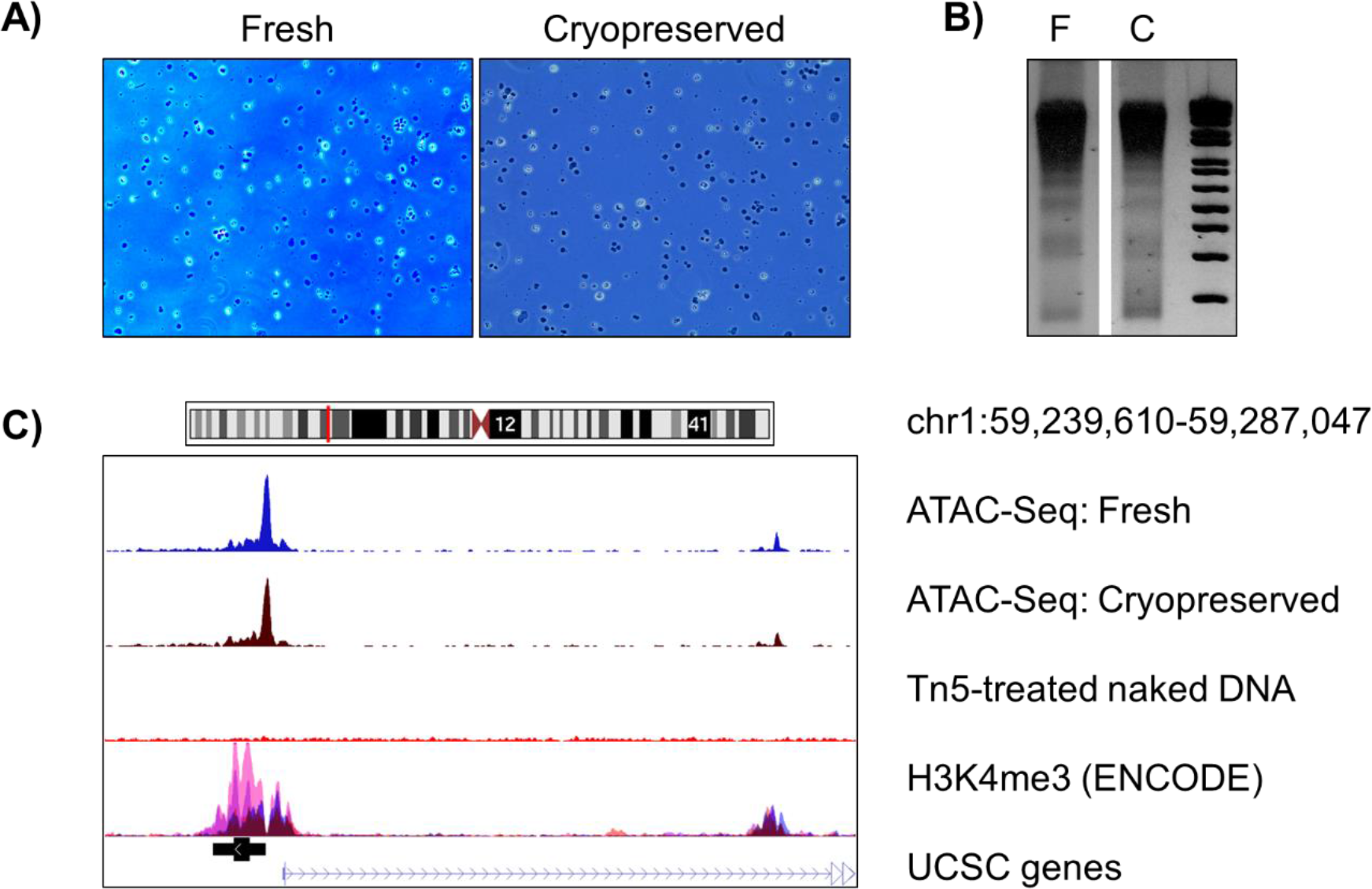
Representative results for ATAC-Seq carried out on fresh and cryopreserved cells. (A) Nuclear morphological evaluation: similar to nuclei from fresh cells, nuclei from cryopreserved neurons were intact and of high quality. (B) Agarose gel electrophoresis of libraries: the nucleosome pattern on the gel was evident for both fresh and cryopreserved samples. (C) ATAC-Seq tracks uploaded on UCSC Genome Browser: peaks from both fresh and cryopreserved neurons were sharp and overlapped with H3K4me3 ChIP-Seq peaks from ENCODE (F = fresh, C = cryopreserved).

**Figure 5.**
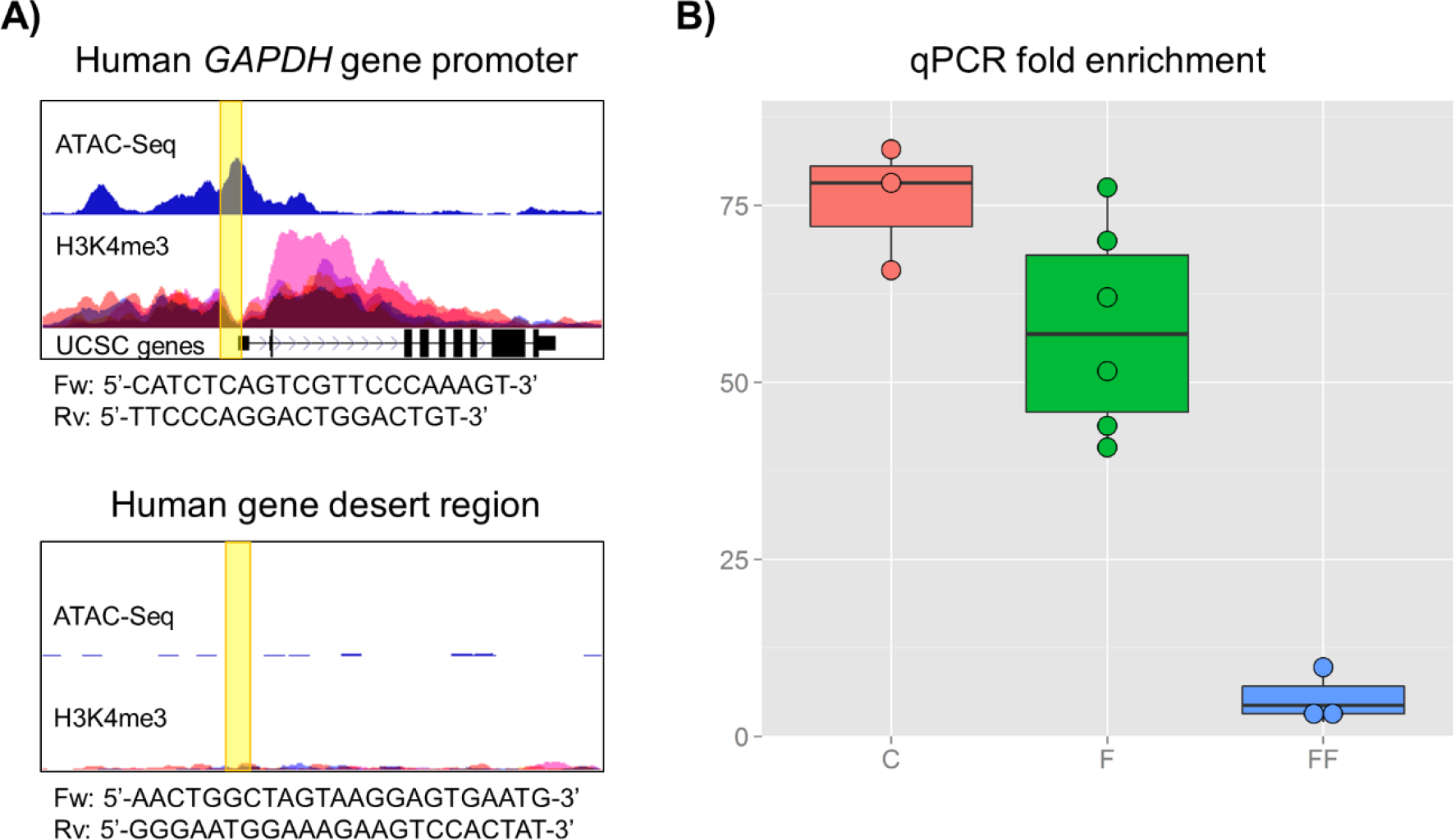
Real-time qPCR for the assessment of the quality of ATAC-Seq libraries. (A) Sequences and genomic locations, displayed on the UCSC Genome Browser, of the primers used to amplify positive and negative control sites. (B) Fold enrichment of the open-chromatin site over the Tn5-insensitive site: while real-time qPCR experiments showed high enrichment for fresh and cryopreserved samples, poor results were obtained with flash-frozen cells (F = fresh, FF = flash-frozen, C = cryopreserved).

### Quantitative comparison of fresh and cryopreserved iMNs

We subsequently performed a series of analyses to quantitatively compare the results from fresh and cryopreserved neurons. We generated sequencing data on three technical replicates from both conditions to assess whether the cryopreservation method induces any modifications in chromatin accessibility. All replicates originated from the same initial batch of cells. Information about sequencing data for each sample is reported in Table 1. The percentage of reads mapping to the human genome was similar for all replicates, but cryopreserved samples displayed higher number of reads mapping to mitochondrial DNA. Despite this discrepancy, we proceeded with our analysis to assess the reproducibility of the epigenetic signal from nuclear DNA across all replicates. To this purpose, we removed mtDNA reads, normalized the libraries to have the same total read counts, and examined the number of reads in 5kb genome windows (excluding ENCODE blacklisted regions). Overall, we observed high reproducibility rates (R ≥ 0.978) between technical replicates in both fresh and cryopreserved samples (Figure 6A). Remarkably, cryopreserved and fresh samples were almost as highly correlated to each other (R ≥ 0.973) as the technical replicates, which suggests that cryopreservation successfully preserves the read distribution across the genome. To further evaluate the similarity between cryopreserved and fresh samples, we identified the peaks in each sample and assigned each one of these peaks to neighboring features (promoters, exons, introns, distal intergenic regions and sites located downstream of the gene) within 1kb (Figure 6B). The distribution of peaks with respect to features in the genome was highly similar across all samples, with most of the peaks located in intergenic regions and promoters. Next, to identify and quantify potential epigenetic alterations induced by the cryopreservation procedure, we performed analysis to detect sites that were significantly different between fresh and cryopreserved samples (see Methods). MACS2-derived peaks across all samples were merged into non-overlapping unique genomic intervals resulting in 75,711 sites. We then used edgeR to detect the differences between the two conditions. We identified very few differentially enriched sites across the genome (210 out of 75,711 total). The magnitude of the differences was small, never exceeding 3-fold (Figure 7). With the exception of chromosome 16, where no differentially enriched sites were identified, the sites spanned the genome, showing no obvious regional biases.

**Figure 6.**
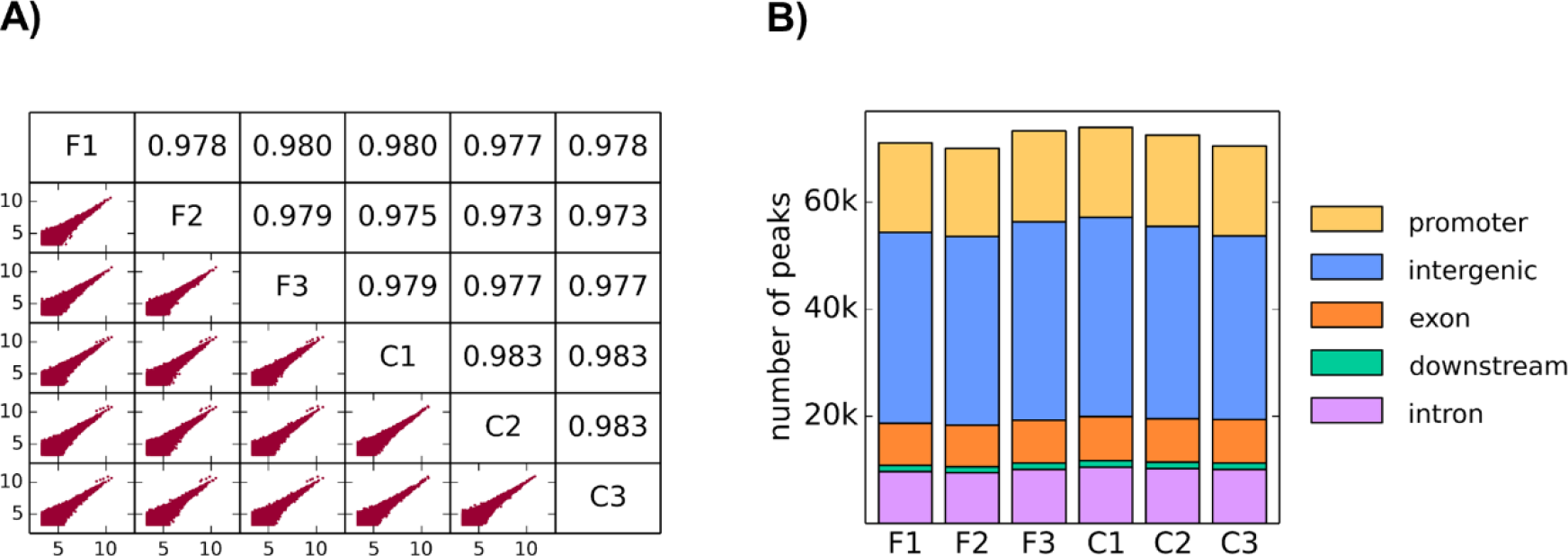
Quantitative comparison of fresh and cryopreserved cells. (A) Correlative analysis of the number of reads in 5kb regions of the genome. The lower left triangle of the figure shows the scatter plots of the log2 read counts for each pair of technical replicates (5kb regions with less than 10 read counts were excluded from the analysis). The upper right triangle displays the corresponding values of the Pearson correlation coefficient. (B) Location-based distribution analysis: the distribution of neighboring genomic features to open-chromatin sites is highly similar between fresh and cryopreserved samples.

**Figure 7.**
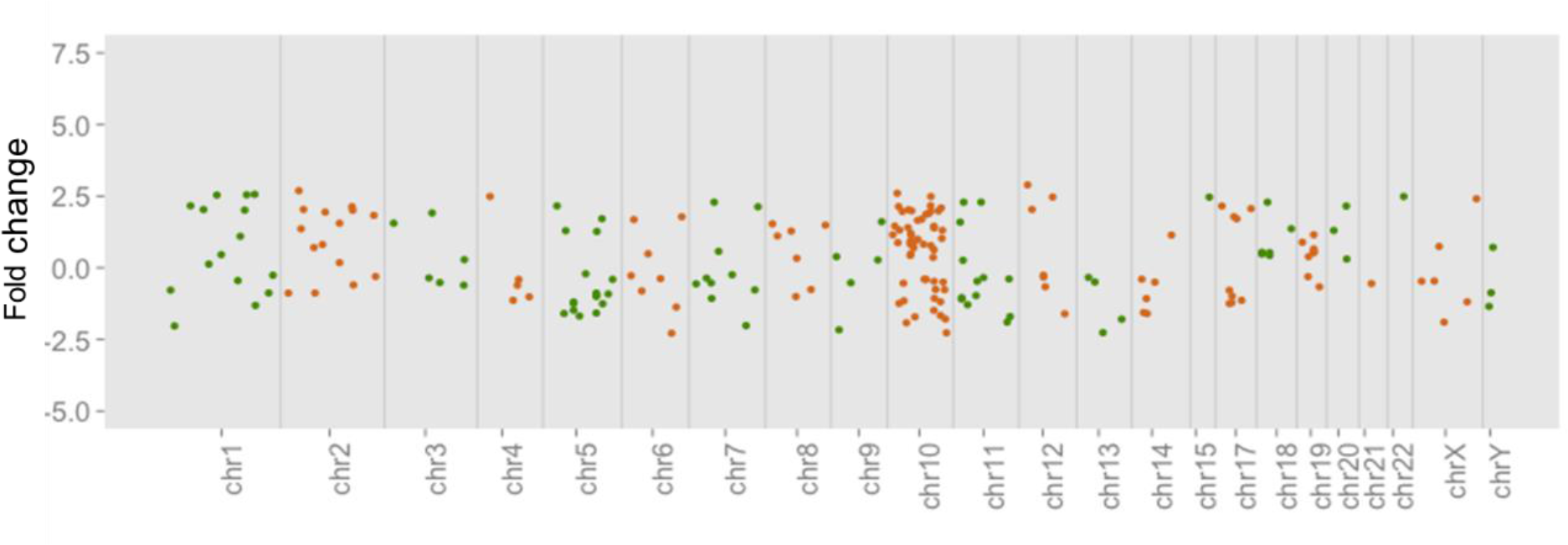
Differentially enriched sites detected between fresh and cryopreserved samples. The fold-change values for differentially enriched peaks between cryopreserved and fresh samples are plotted as a function of the position of the sites across all genome. The changes were small (< 3-fold) and no regional bias for these sites was observed.

In conclusion, we established a cell freezing protocol suitable for ATAC-Seq experiments on iPSC-derived motor neurons. As in the case of fresh samples, the cryopreserved samples passed all of the quality control checkpoints. Although we observed that higher numbers of reads map to mitochondria DNA in cryopreserved iMNs, we demonstrated that the epigenetic signal from nuclear DNA was highly reproducible between fresh and cryopreserved neurons. This optimized protocol provides a simple and practical approach to extend ATAC-Seq to frozen cells. We anticipate that this work will be of great value to epigenetic investigators.

## Methods

### Primary cells and iPSC derivation

Source fibroblast lines were obtained from Coriell (GM09677) under institutional review board approved protocols. The fibroblast-derived iPSC line 77iSMA-n5 was created by the Cedars-Sinai Medical Center iPSC Core using the episomal vectors pCXLE-hUL, pCXLE-hSK, and pCXLE-hOCT3/4-shp53-F (Addgene, from a previously published protocol^8^). We transfected the fibroblasts with the vectors using the Amaxa Human Dermal Fibroblast Nucleofector Kit. The 77iSMA-n5 line was characterized by the Cedars-Sinai iPSC Core using the following quality control assays: G-Band karyotyping, immunocytochemistry for pluripotency markers, embryoid body formation, PluriTest, and qRT-PCR for endogenous pluripotency genes^8-12^.

### Motor Neuron Precursors (iMPs)

The SMA patient line, 77iSMA-n5, was grown until 90% confluent using a standard iPSC maintenance protocol. On Day 0 of differentiation, iPSCs were lifted as single cells by Accutase treatment for 5 min at 37°C. We counted the cells and re-suspended them in Neuroectoderm differentiation media (NDM+LS), which contains 1:1 IMDM/F12, 1% NEAA, 2% B-27, 1% N2, 1% Antibiotic-Antimycotic, 0.2 μΜ LDN193189 and 10 μΜ SB431542. Next, we seeded 25,000 cells/well in a 384-well plate and centrifuged the cells for 5 min at 200 rcf. On day 2, we transferred the neural aggregates to a poly 2-hydroxyethyl methacrylate (poly-Hema) coated flask and cultured them for an additional 3 days in NDM+LS media. On day 5, we seeded the neural aggregates onto a tissue culture plate coated with laminin (50μg/mL) to induce rosette formation. From day 12-18, the attached neural aggregates were transitioned to Motor Neuron Specification Media (1:1 IMDM/F12, 1% NEAA, 2% B-27, 1% N2, 1% Antibiotic-Antimycotic, 0.25 μM all-trans retinoic acid (ATRA), 1 μM purmorphamine (PMN), 20 ng/mL brain-derived neurotrophic factor (BDNF), 20 ng/mL glial cell line-derived neurotrophic factor (GDNF), 200 ng/mL ascorbic acid (AA) and 1 μM dibutyryl cyclic-AMP (db-cAMP). On day 19 we selected the rosettes by incubating them with Neural Rosette Selection Reagent (StemCell Technologies Cat#05832) for 45 min at 37°C. After selection, we collected the rosettes and transferred them to poly-Hema coated T75 flasks and cultured the cells as iMPs in Motor Neuron precursor expansion media (MNPEM), which contains 1:1 IMDM/F12, 1% NEAA, 2% B27, 1% N2, 1% Antibiotic-Antimycotic, 0.1μΜ ATRA, 1 μΜ PMN, 100ng/mL EGF and 100ng/mL FGF2. We expanded the iMPs as aggregates in suspension using a mechanical passaging method known as “chopping”^13,14^ for up to five passages. For cryopreservation, we pooled the aggregates and dissociated them via a combined enzymatic (Accutase for 10 minutes at 37°C) and mechanical dissociation strategy to form a single cell suspension. The single cell suspension was then concentrated via centrifugation (200 rcf for 5 minutes at 4°C), re-suspended in Cryostor (StemCell Technologies Cat #: 07930), cryopreserved using a controlled rate freezer (Planer Inc.) and stored in gas-phase liquid nitrogen.

### Motor Neuron Cultures (iMNs)

We derived the iMNs by thawing the iMPs and immediately plating the single cell suspension onto plastic tissue culture-treated plates coated with 50μg/mL laminin for two hours at 37°C. We seeded the iMPs in Motor Neuron Maturation Medium (MNMM) Stage 1 consisting of 1:1 IMDM/F12, 1% NEAA, 2% B-27, 1% N2, 1% Antibiotic-Antimycotic, 0.1 μM ATRA, 1 μM PMN, 10 ng/mL BDNF, 10 ng/mL GDNF, 200 ng/mL AA, 1 μM db-cAMP, and 2.5 μM N-[(3,5-Difluorophenyl)acetyl]-L-alanyl-2-phenyl]glycine-1,1-dimethylethyl ester (DAPT). We cultured the cells for a period of seven days. On day 7, the plated cultures were transitioned to MNMM Stage 2 containing 98.8% Neurobasal media, 1% non-essential amino acids, 0.5% Glutamax, 1% N2, 10 ng/mL BDNF, 10 ng/mL GDNF, 200 ng/mL AA, 1 μM db-cAMP, and 0.1 μM Ara-C. We further differentiated the iMNs in MNMM Stage 2 for a total of 21 days. On day 21, the iMNs cultures were either fixed for immunocytochemistry or collected.

### Cell collection, freezing and thawing

For cell collection, the iMNs were washed once with 1X PBS, isolated via cell scraper in 1X PBS, and centrifuged at 200 rcf for 5 minutes at 4°C.

Flash-freezing: pellets (no supernatant) were flash-frozen in liquid nitrogen.

Cryopreservation: pellets were re-suspended in Cryostor media and frozen slowly in a Mr. Frosty isopropyl alcohol chamber (FisherSci) at −80°C overnight. This procedure allowed us to achieve a rate of cooling of −1°C/minute.

Both the flash-frozen isolated cell pellets and the cryopreserved iMNs were stored at −80°C until experimentation. To thaw the cryopreserved iMNs, we removed the cryovials from −80°C and quickly warmed them for 2 min in a 37°C water bath. We transferred the samples to 12 ml of warm 1X PBS supplemented with 1X protease inhibitor cocktail. We gently mixed each tube by inversion and centrifuged at 200 rcf for 5 min at 4°C. We carefully aspirated the supernatant and proceeded with nuclei isolation. Flash-frozen cell pellets were removed from −80°C and immediately re-suspended in ice-cold cell lysis buffer.

### Immunocytochemistry

We fixed iMNs with 4% paraformaldehyde and blocked them with 5% donkey serum with 0.1% Triton X-100 in 1X PBS. We incubated the cells overnight at 4°C with the following primary antibodies: anti-SMI32 (mouse monoclonal, 1:1000, BioLegend, cat. no. SMI-32R) and anti-ISL1 (goat polyclonal, 1:250, R&D Systems, cat. no. AF1837). We subsequently rinsed the cells and incubated them with species-specific Alexa Fluor 488-conjugated secondary antibody (donkey anti-mouse immunoglobulin G (IgG), 1:1000, Life Technologies, cat. no. A-21202) and Alexa Fluor 594-conjugated secondary antibody (donkey anti-goat IgG, 1:1000, Life Technologies, cat. no. A-11058). We counterstained nuclei using DAPI (1 μg/mL). We acquired the images using Nikon/Leica microscopes with 10x magnification.

### Purification of nuclei from iMNs

We re-suspended the cell pellets in ice-cold cell lysis buffer (10 mM Tris-HCl, pH 7.4, 10 mM NaCl, 3 mM MgCl2, 0.1% IGEPAL CA-630) supplemented with 1X protease inhibitor cocktail (Roche). We incubated the cells on ice for 5 min and centrifuged at 230 rcf for 5 min at 4°C. We carefully removed the supernatant and re-suspended the nuclei in 25 μl of ice-cold 1X Tagment DNA Buffer (Illumina). We quantified the nuclei with Trypan Blue staining and the Countess^®^ Automated Cell Counter (Invitrogen).

### DNA extraction

We purified the DNA from iMNs using the DNeasy Blood & Tissue Kit (Qiagen), according to the manufacturer's instructions. We quantified the DNA using a NanoDrop 2000 instrument (Thermo Scientific) and used 50 ng to prepare the DNA library using the Nextera DNA Library Preparation Kit (Illumina), according to the manufacturer's instructions. This library, obtained from naked DNA, was used as internal control to determine the background level of intrinsic accessibility of genomic DNA and correct for any Tn5 transposase sequence cleavage bias.

### Chromatin tagmentation and sequencing

We used 50,000 nuclei for the transposase reaction, which was carried out as described in Buenrostro et al.^1^. We subsequently purified the samples with the DNA Clean & Concentrator-5 Kit (Zymo Research) and eluted them with 20 μl of Elution Buffer (Qiagen). We PCR-amplified the samples using 25 μl of Nextera PCR Master Mix (Illumina), 5 μl of PCR Primer Cocktail (Illumina), 5 μl of Index primer 1 (i7, Illumina), and 5 μl of Index primer 2 (i5, Illumina). We used the following PCR reaction protocol: 3 min 72°C; 30 sec 98°C; 8 cycles (10 sec 98°C, 30 sec 63°C, 3 min 72°C). We purified the samples with the DNA Clean & Concentrator-5 Kit (Zymo Research), eluted them with 20 μl of Elution Buffer (Qiagen), and loaded them on 2% agarose gel (Invitrogen) for qualitative evaluation of libraries and size-selection. We size-selected the following fractions: 175 - 250 bp (fraction “A”, corresponding to a nucleosome-free fragment size) and 250 - 625 bp (fraction “B”). We purified the DNA from both gel fractions, using the QIAquick Gel Extraction Kit (Qiagen) following the manufacturer’s recommendation, and eluted it with 20 μl of Elution Buffer (Qiagen). We utilized the DNA from fraction “B” for qPCR-based qualitative analysis of libraries using primers mapping to open-chromatin regions as positive control sites and gene desert regions as negative control sites (Figure 5). We also performed the qPCR assay using 10-fold serial dilutions of non-transposed genomic DNA as a template to generate a calibration line for each primer pair and correct for any differences in the primer efficiency. The fold enrichment of the open-chromatin site (OC) over the Tn5-insensitive site (INS) was calculated with the following formula, as previously described^15^: 2 to the power of [(OCn-OCa) - (INSn-INSa)], where OCn is the qPCR threshold cycle number obtained for the OC qPCR primer pair using transposed naked DNA as template, and INSa is the qPCR threshold cycle number obtained for the INS qPCR primer pair using ATAC-Seq library as template. As an additional control, we carried out the qPCR assay using transposed naked DNA. No fold-enrichment of open-chromatin sites should be detected when using transposed naked DNA as a template. We prepared the amplification reaction with 1X KAPA SYBR FAST qPCR Master Mix (Kapa Biosystems) and 500 nM of forward and reverse primers. We carried out qPCR assays using a LightCycler^®^ 480 Instrument II (Roche), available at the MIT BioMicroCenter. We further amplified the DNA from fraction “A” with 1X NEBNext High-Fidelity PCR Master Mix (New England Biolabs), 200 nM of Primer 1 (5'-AATGATACGGCGACCACCGA-3'), and 200 nM of Primer 2(5'-CAAGCAGAAGACGGCATACGA-3’). We used the following PCR reaction protocol: 30 sec 98°C; 4 cycles (10 sec 98°C, 30 sec 65°C, 30 sec 72°C); 5 min 72°C. We purified the final libraries using Agencourt AMPure XP beads (Beckman Coulter), checked their quality using a Fragment Analyzer™ instrument (Advanced Analytical), and measured their concentration by a qPCR-based method (KAPA Library Quantification Kit for Illumina Sequencing Platforms). We submitted the samples to the MIT BioMicroCenter for single-end sequencing with the Illumina HiSeq 2000 platform.

### Bioinformatic analysis

We aligned sequencing reads to the hg19 genome build using BWA. We assessed the quality of the sequences using FastQC (more details on how the data was processed can be found at http://openwetware.org/wiki/BioMicroCenter:Software#BMC-BCCPipeline). Given the large percentage of mitochondrial reads found in some samples, we removed mitochondrial reads from the analysis using custom UNIX scripts. We determined open-chromatin regions (peaks) using MACS^16^ v.2.1.0.20150420 (q-value threshold = 0.05). We used the sequencing data from transposed naked DNA as a control. The differential analysis was performed using the default settings in the package DiffBind^17^ version 1.16.0. Briefly, read counts for each site were computed and differentially enriched sites between fresh and cryopreserved conditions were identified using the edgeR package^18^, with FDR < 0.1.

## Acknowledgments

This work was supported by the National Institute of Health (U54-NS-091046 and U01-CA-184898), and used computing resources funded by the National Science Foundation (DB1-0821391) and sequencing support from the National Institutes of Health (P30-ES002109) through the MIT BioMicro Center. The authors are particularly grateful to Dr. Andrea Allais for his assistance with the figures and valuable comments on the manuscript.

## Author contributions

P.M., C.N.S. and E.F. designed the study. P.M., R.E., B.C.S., N.P. and E.F. wrote the manuscript. P.M. carried out ATAC-Seq experiments. N.P., X.X. and M.A. assisted with protocol optimization and experiments. P.M. and R.E. performed the analysis. B.C.S, B.M. and D.S. generated iPSCs, iMPs and iMNs. All authors reviewed the manuscript.

